# Brain glucose metabolism and ageing: A 5-year longitudinal study in a large PET cohort

**DOI:** 10.1101/2022.09.15.508088

**Authors:** Kyoungjune Pak, Tuulia Malén, Severi Santavirta, Seunghyeon Shin, Hyun-Yeol Nam, Sven De Maeyer, Lauri Nummenmaa

## Abstract

**Background:** Ageing and clinical factors impact brain glucose metabolism. However, there is a substantial variation of the reported effects on brain glucose metabolism across studies due to the limited statistical power and cross-sectional study designs.

**Methods:** We retrospectively analyzed data from 441 healthy males (mean 42.8, range 38-50 years) who underwent health check-up program twice at baseline and 5-year follow-up. Health check-up program included 1) brain ^18^F-Fluorodeoxyglucose (FDG) positron emission tomography (PET), 2) anthropometric and body composition measurements, 3) blood samples, and 4) questionnaires for stress and depression. After spatial normalization of brain FDG PET scans, standardized uptake value ratio (SUVR) was measured from 12 region-of-interests. We used hierarchical clustering analysis to reduce their dimensionality before the Bayesian hierarchical modelling. Five clusters were established for predicting regional SUVR; 1) metabolic cluster (body mass index, waist-to-hip ratio, fat percentage, muscle percentage, homeostatic model assessment index-insulin resistance), 2) blood pressure (systolic, diastolic), 3) glucose (fasting plasma glucose level, HbA1c), 4): psychological cluster (stress, depression), and 5) heart rate. The effects of clinical variable clusters on regional SUVR were investigated using Bayesian hierarchical modelling with brms that applies the Markov-Chain Monte Carlo sampling tools.

**Results:** All the clinical variables except depression changed during the 5-year follow-up. SUVR decreased in caudate, cingulate, frontal lobe and parietal lobe and increased in cerebellum, hippocampus, occipital lobe, pallidum, putamen, temporal lobe and thalamus. SUVRs of thalamus, pallidum, hippocampus, putamen and parietal lobe were negatively associated with metabolic cluster and the effects of glucose on SUVRs varied across regions. SUVRs of thalamus, hippocampus, cingulate, cerebellum increased and those with occipital lobe decreased with heart rate. The effects of blood pressure and psychological cluster markedly overlapped with zero across regions.

**Conclusion:** Regionally selective decline in brain glucose utilization begins already in the middle age, while individual differences in brain glucose metabolism remain stable. In addition to ageing, brain glucose utilization is also associated with metabolic cluster, blood glucose levels and heart rate. These effects are also consistent over the studied period of 5 years in the middle adulthood.

## INTRODUCTION

Human brain utilizes glucose as its main source of energy, thus, brain glucose metabolism, assessed by positron emission tomography (PET) with ^18^F-Fluorodeoxyglucose (FDG) can be utilized for quantifying neuronal activity in the human brain (1). Ageing is accompanied by structural changes of brain including overall shrinkage of brain volume and cortical thinning (2), alterations in the functioning of specific neuroreceptor systems (3; 4) and decrease in brain glucose metabolism in frontal, temporal, parietal lobes (5; 6). Understanding the age-dependent alterations in cerebral glucose utilization is imperative, given that ageing and concomitant atrophy and metabolic changes in the brain may lead to decline in cognitive functions including memory (2), executive functions such as planning and inhibitory control (7).

However, there is substantial variation in the age-dependent effects on brain glucose metabolism across studies, and this variation pertains both the direction of the effect as well the regions affected by ageing (8–10). It is possible this variation results from the between-subjects design in the studies and lacking statistical power, leading to false positive and negative findings (11–13). Only a few longitudinal studies on brain glucose metabolism have been conducted, focusing on mainly Alzheimer’s dementia (14–16). The annual decrease of brain glucose metabolism in Alzheimer’s dementia-related regions such as precuneus/posterior cingulate, lateral parietal/temporal cortices was 0.22% in healthy subjects (14). However, as a small number (n=107) of older age adults (mean 67.9 years) was included in the study by Ishibashi et al (14), there is a limitation to define the ageing effects in middle-aged adults and longitudinal studies of healthy middle-aged adults would be necessary.

In addition to ageing, metabolic variables have effects on brain glucose uptake. Body mass index (BMI) is negatively associated with brain glucose metabolism in prefrontal cortex and cingulate gyrus (17), but some studies have found a positive association in orbitofrontal cortex for females (18). Skeletal muscle area is positively associated with a lower probability of Alzheimer's dementia assessed by ^18^F-FDG PET (19). Also, homeostatic model assessment index-insulin resistance (HOMA-IR), indexing insulin resistance, is negatively associated with brain glucose metabolism in frontal, parietal and temporal lobes (20). Finally, several studies have reported a negative association between blood glucose level and brain glucose metabolism of posterior cingulate gyrus (21–23), occipital cortex (22) or prefrontal cortex (23), with the inconsistency of regions across the studies. However, a study using euglycemic hyperinsulinemic clamp showed insulin sensitivity, indexed by M value was negatively associated with brain glucose metabolism in all brain regions, in the opposite way compared with skeletal muscle (5).

### The current study

There is evidence that both ageing and metabolic factors influence brain glucose metabolism, but the effects have been inconsistent across studies, possibly due to the limited statistical power and cross-sectional study designs. To address the effects of age, metabolic and psychological factors on brain glucose utilization, we analyzed a large cohort (n=441) of healthy middle-aged adults (mean 42.8 years) who underwent ^18^F-FDG PET scans and a health check-up program twice: at the baseline and at 5-year follow-up. We used Bayesian hierarchical modeling to estimate the effects of clinical variables on brain glucose metabolism and show that ageing and metabolic risk factors are negatively associated with brain glucose metabolism, while blood glucose levels have a regionally variable positive and negative effects.

## MATERIALS AND METHODS

### Subjects

We retrospectively analyzed data from 473 healthy males who underwent health check-up program at Samsung Changwon Hospital Health Promotion Center in 2013 (baseline) and 2018 (follow-up). After excluding subjects with neuropsychiatric disorders (n=5) or malignancies (n=3), those with missing data of anthropometric and body composition measurements (n=24), 441 healthy males were included in both baseline (mean 42.8, range 38-50 years) and follow-up (mean 47.9, range 43-55 years) studies. Health check-up program included 1) ^18^F-FDG PET, 2) anthropometric and body composition measurements, 3) blood samples, and 4) questionnaires of stress and depression. The study protocol was approved by the Institutional Review Board and the informed consent from the participants was waived due to the retrospective study design.

### ^18^F-FDG PET

Subjects were asked to avoid strenuous exercise for 24 hours and fast for at least 6 hours before PET study. PET/computed tomography (CT) was performed 60 mins after injection of ^18^F-FDG (3.7 MBq/kg) with Discovery 710 PET/CT scanner (GE Healthcare, Waukesha, WI, USA). Continuous spiral CT was obtained with a tube voltage of 120kVp and tube current of 30-180mAs. PET scan was obtained in 3-dimensional mode with full width at half maximum of 5.6 mm and reconstructed using an ordered-subset expectation maximization algorithm. PET scans were spatially normalized to MNI space using PET templates from SPM5 (University College of London, UK) with pmod version 3.6 (PMOD Technologies LLC, Zurich, Switzerland). Automated Anatomical Labeling 2 (AAL2) atlas (24) was used to define region-of-interests (ROIs); caudate, cerebellum, cingulate, frontal lobe, hippocampus, insula, occipital lobe, pallidum, parietal lobe, putamen, temporal lobe and thalamus. The mean uptake of each ROI was scaled to the mean of global cortical uptake of each individual, and defined as standardized uptake value ratio (SUVR). Percentage change of SUVR from each region was calculated as follows: (follow-up SUVR–baseline SUVR)/baseline SUVR*100 (%). To compare baseline and follow-up scans of each subject, paired t-test was computed using SPM12 after smoothing the SUVR images with a Gaussian kernel of FWHM 8mm.

### Body measurement

For anthropometric measurements, height (cm) and weight (kg) were measured and body mass index (BMI) was calculated as weight/height^2^ (kg/m^2^). Waist and hip circumference (cm) were measured, and wait-to-hip ratio was calculated. For body composition measurement, a commercially available bioelectrical impedance analysis (InBody S10, Biospace, Seoul, Republic of Korea) was used to calculate fat percentage (%) and muscle percentage (%) after dividing fat mass (kg) and skeletal muscle mass (kg) with weight (kg). Systolic and diastolic blood pressures (mmHg) and heart rate were measured using an automatic sphygmomanometer (EASY X 800; Jawon Medical Co., Ltd, Seoul, Korea) after at least 10 mins of rest.

### Blood samples

Blood samples were collected from the antecubital vein of each subject. Fasting plasma glucose (mg/dL) and insulin (μIU/mL) were measured and HOMA-IR was calculated as follows: fasting insulin (μIU/mL)*fasting plasma glucose (mg/dL)/405 (25). Hemoglobin A1c (HbA1c, %) levels were analyzed with high-performance liquid chromatography.

### Questionnaires

Subjects completed a standardized questionnaire battery on stress and depression. Stress was measured with the stress questionnaire developed for the Korea National Health and Nutrition Examination Survey (26). It consists of 9 items on a scale of 1 to 5, regarding the experienced stress over the previous month. Depression was measured with the Centre for Epidemiologic Studies-Depression (Korean version), a self-report questionnaire designed to measure depressive symptoms over the previous week, including 20 items on a scale of 0 to 3 (27).

### Statistical analysis

Normality was tested with Shapiro-Wilk test and comparison of clinical variables and SUVRs between baseline and follow-up was assessed with Wilcoxon matched-pairs signed rank test.

### Cluster analysis for predictor variables

Because some clinical variables contained missing values (baseline study: HOMA-IR 41/441, 9.3%, stress score 15/441, 3.4%, depression score 31/441, 7.0%; follow-up study: questionnaire - stress 3/441, 0.7%, depression 4/441, 0.9%), we used a non-parametric imputation algorithm missForest with default parameters for imputation of the missing values (28). As some clinical variables were strongly correlated, we used hierarchical clustering analysis before the Bayesian hierarchical modelling. Before clustering, muscle percentage was multiplied with −1 to simplify the solution as it was the only variable showing negative correlations with the other predictors. Clustering yielded stable cluster hierarchy across the 6 tested algorithms (complete-linkage, single-linkage, UPGMA, WPGMA, WPGMC and Ward) and the following clusters were defined 1) metabolic cluster (BMI, waist-to-hip ratio, fat percentage, muscle percentage, HOMA-IR), 2) blood pressure (systolic, diastolic), 3) glucose (fasting plasma glucose level, HbA1c), 4): psychological cluster (stress, depression), and 5) heart rate. In each cluster, the value was calculated after averaging the standardization of clinical variables.

### Bayesian hierarchical modelling

The effects of clinical clusters on regional SUVR were investigated using Bayesian hierarchical modelling with brms (29–31) that applies the Markov-Chain Monte Carlo sampling tools of RStan (32). We set up a model with regional SUVR as a dependent variable and 5 clusters (metabolic, blood pressure, glucose, psychological, and heart rate) as predictors. These fixed effects were calculated individually and as an interaction with time. We added subject and ROI as random intercepts to allow SUVR to vary between subjects and ROIs and calculated the fixed effects including the interactions with time separately for each ROI as a random slope. Bayesian models were estimated using four Markov chains, each of which had 4,000 iterations including 1,000 warm-ups, thus totaling 12,000 post-warmup samples. The sampling parameters were slightly modified to facilitate convergence (max treedepth = 20). Statistical analysis was carried out in R Statistical Software (The R Foundation for Statistical Computing).

## RESULTS

Mean SUVRs for baseline and follow-up ^18^F-FDG PET scans are shown in Figure 1 and subjects’ characteristics are summarized in Table 1. Except for depression score (p=0.3332) all the clinical variables changed during the 5-year follow-up. Stress score (p<0.0001) and muscle percentage (p<0.0001) decreased while other variables (BMI, waist-hip ratio, fat percentage, HOMA-IR, systolic and diastolic blood pressure, glucose, HbA1c, and heart rate) increased (ps<0.0001).

**Figure 1.**
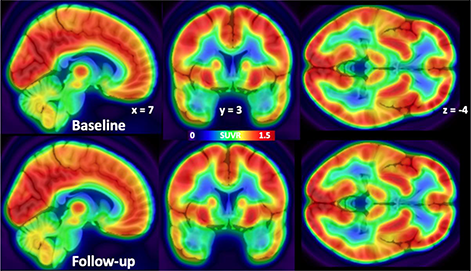
Mean SUVR map of ^18^F-FDG PET scans from 441 subjects

**Table 1.**
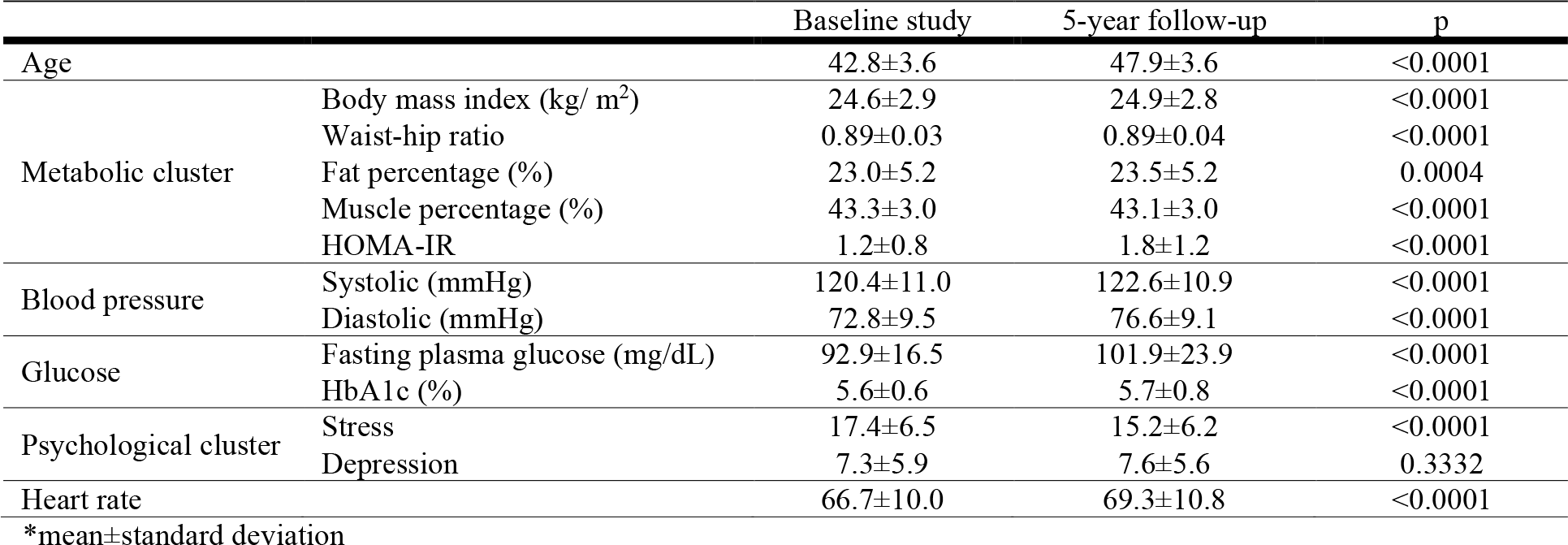
Subjects’ characteristics

SUVRs were correlated across the baseline and follow-up scans (Figure 2, mean correlation coefficient of 0.8645). Age at baseline was negatively correlated with SUVR in insula (rho=−0.1375, p=0.0038) but not in other brain regions.

**Figure 2.**
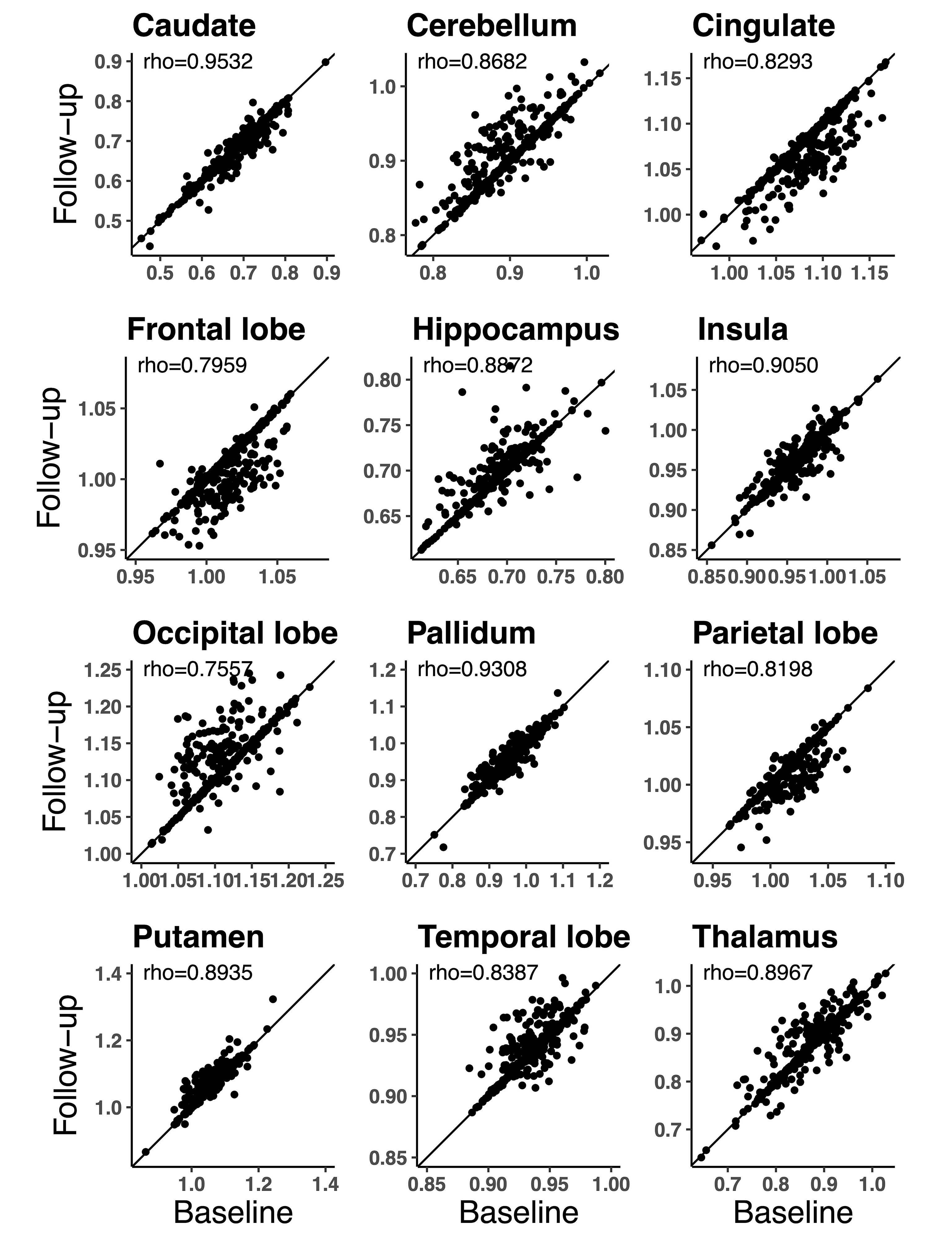
Distribution of SUVRs from 12 ROIs in baseline and follow-up ^18^F-FDG PET scans

During the 5-year follow-up period, SUVR decreased in caudate (p<0.0001), cingulate (p<0.0001), frontal lobe (p<0.0001 and parietal lobe (p<0.0001) and increased in cerebellum (p<0.0001), hippocampus (p<0.0001), occipital lobe (p<0.0001), pallidum (p=0.0479), putamen (p<0.0001), temporal lobe (p=0.0310) and thalamus (p=0.0236). SUVR in insula (p=0.6975) remained unaltered (Figure 2). Mean percentage changes of SUVRs during 5-year follow-up were −0.4712 % (caudate), 0.9642 % (cerebellum), −0.7843 % (cingulate), −0.5350 % (frontal lobe), 0.5543 % (hippocampus), −0.0297 % (insula), 1.0201 % (occipital lobe), 0.2224 % (pallidum), −0.4077 % (parietal lobe), 0.7020 % (putamen), 0.2389 % (temporal lobe) and 0.5439 % (thalamus), respectively. Complementary full volume comparison between baseline and follow-up SUVR is shown in Figure 3; the results accord with the ROI-based analysis.

**Figure 3.**
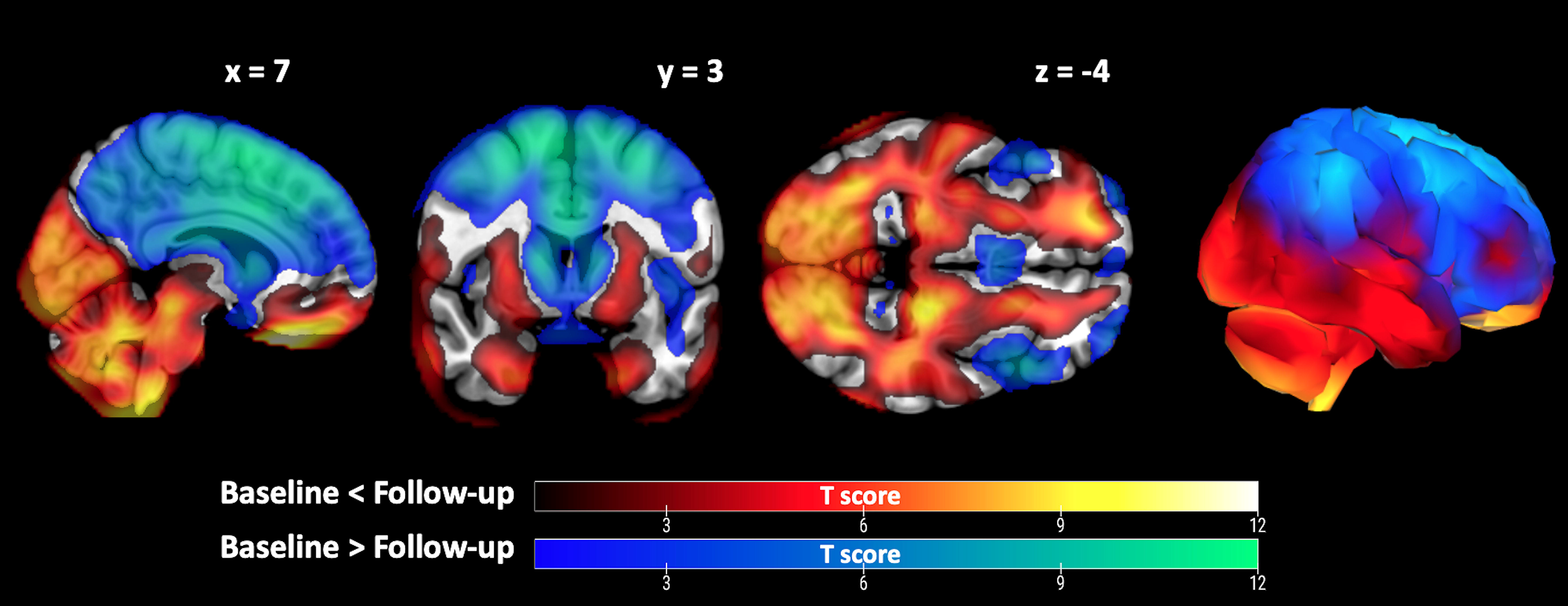
Full volume comparison between baseline and follow-up ^18^F-FDG PET scans

The effects of clinical variables on SUVRs were generally similar in the baseline and follow-up measurements. SUVRs of thalamus, pallidum, hippocampus, putamen and parietal lobe were negatively associated with metabolic cluster, whereas that of frontal lobe was positively associated with metabolic cluster. The effects of glucose on SUVRs varied across regions. Glucose of baseline and follow-up studies was positively associated with SUVRs of hippocampus, caudate, temporal lobe, cerebellum and negatively associated with those of parietal lobe, cingulate, occipital lobe, frontal lobe, some of their 95% posterior intervals overlapping with zero. The effects of blood pressure and psychological cluster markedly overlapped with zero across regions, except for negative association between SUVR of thalamus, hippocampus from follow-up ^18^F-FDG PET scan and psychological cluster. SUVRs of thalamus, hippocampus, cingulate, cerebellum increased and those with occipital lobe decreased with heart rate, some of their 95% posterior intervals overlapping with zero (Figure 4).

**Figure 4.**
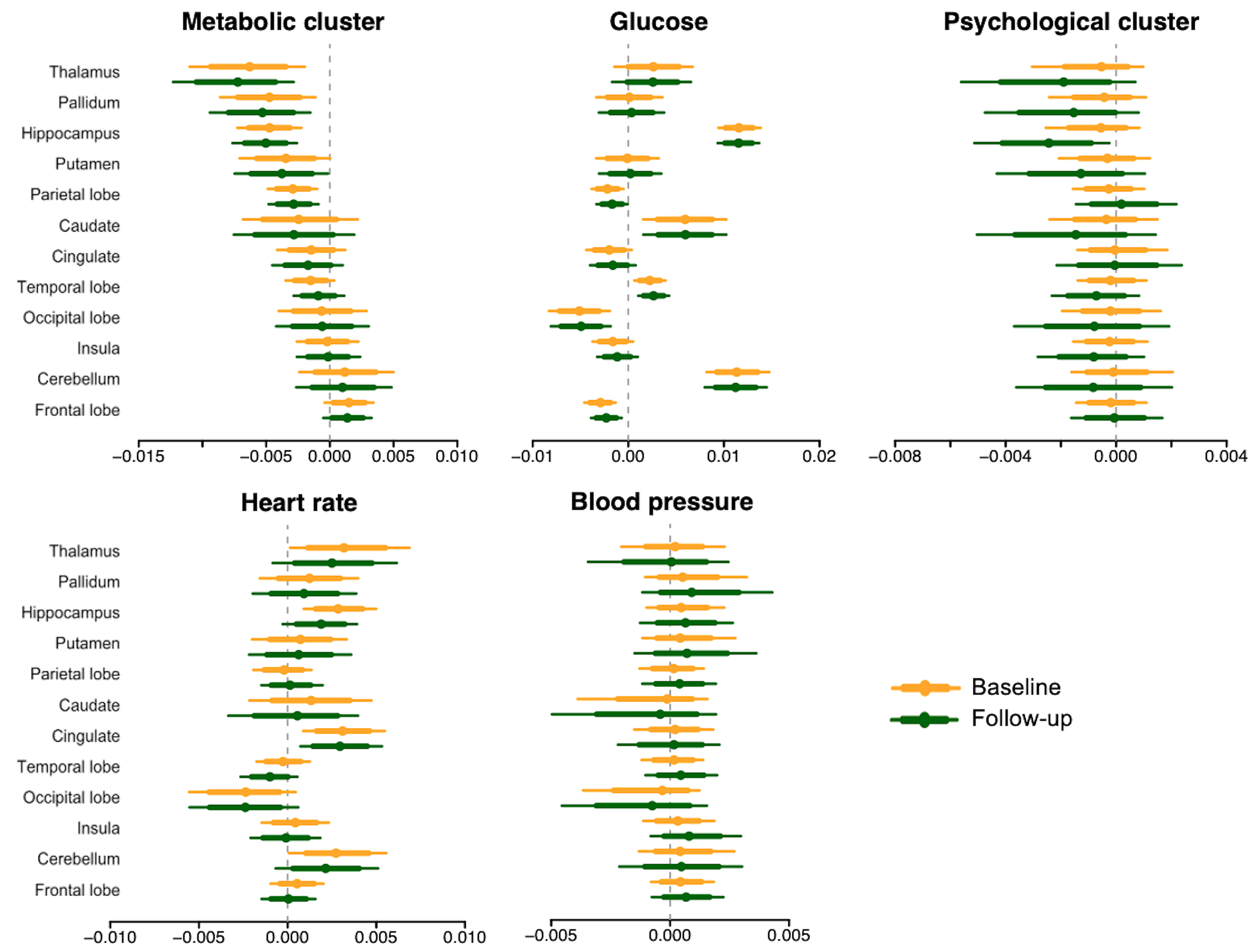
Posterior intervals of the regression coefficients for each cluster predicting SUVR. The thick lines represent the 80% posterior intervals, the thin lines represent the 95% posterior intervals, and the circles represent posterior means

## DISCUSSION

Our main finding was that ageing led to decrease in brain glucose metabolism in caudate, cingulate, frontal lobe, and parietal lobe, while increased brain glucose metabolism was observed in cerebellum, hippocampus, occipital lobe, pallidum, putamen, temporal lobe, and thalamus.

Additionally, metabolic variables, blood glucose levels and heart rate had regionally specific associations with brain glucose metabolism. These effects also remained consistent over the 5-year follow-up, and the regional brain glucose metabolism were strongly correlated between the baseline and follow-up studies.

### Ageing effect

The brain uses glucose as its principal substrate for energy synthesis under normal physiological conditions. Therefore, it requires a constant supply of glucose from the bloodstream to maintain the function (33–35). FDG, an analog of glucose, has long been used to measure cerebral metabolic rates of glucose, thus providing a proxy for neuronal activity (33). Glucose utilization as measured with FDG PET represents mostly neuronal and synaptic activities in brain tissue. Currently, FDG PET is the most accurate in vivo method for the investigation of brain glucose metabolism in clinical settings (36).

After 5-year follow-up, brain glucose metabolism of caudate, cingulate, frontal lobe, parietal lobe was decreased and those from cerebellum, hippocampus, occipital lobe, pallidum, putamen, temporal lobe, and thalamus was increased. These effects were observed although the participants were relatively young (baseline mean 42.8, range 38-50 years; follow-up mean 47.9, range 43-55 years). Within this age range the annual decline of brain glucose metabolism was approximately 0.2% or less across the regions, slightly below than what has been reported in previous longitudinal studies of 0.22% to 0.58% for cingulate, parietal and temporal lobe in healthy older adults (mean 67.9 years) (14–16). Glucose utilization decline thus starts already during middle age and is further accelerated in the elderly (37).

During normal ageing, brain function and structure are also altered in the absence of clinically severe deficits (38). Here, we found that SUVR decreased as ageing effects particularly in the frontal lobe. This accords with prior studies where lowered frontal glucose metabolism is the most consistent finding in FDG PET studies of ageing, with typical decline being 12-13 % between 20 and 70 years of age (39; 40). As frontal lobe is involved in executive functions and attentional performance, this is thought to be related with cognitive decline occurring in normal ageing (41). In addition to frontal lobe, SUVRs in caudate, cingulate and parietal lobe were also decreased, consistent with previous studies (6; 15), which might have an association with the enlargement of the cerebroventricular space, the widening of sulci and the decline of gray matter volume (42).

We also found that SUVRs in cerebellum, hippocampus, occipital lobe, pallidum, putamen, temporal lobe and thalamus were increased after 5-year follow-up, consistent with previous studies (10; 41). Functional magnetic resonance imaging (fMRI) studies have found that ageing is associated with increased activity in the visual, motor and subcortical regions as a compensatory response to decreased activity in the default mode network (43). In addition, tau accumulation in the ageing brain might lead to increased glucose metabolism. In normal ageing, higher tau accumulation is associated with increased brain glucose metabolism of ^18^F-FDG PET (44; 45). Even early as the age of 47.4 years, half of the population can be affected by the neurofibrillary pathology, aggregation of hyperphosphorylated tau protein (46). Tau accumulation starts from the entorhinal cortex in temporal lobe, followed by hippocampus, which is consistent with the regions of increased glucose metabolism after 5-year follow-up in this study. Because tau accumulation was not directly investigated in the present study, this needs to be elucidated in future studies.

### Brain glucose metabolism and clinical variables

Metabolic cluster consisting of well-known metabolic risk factors (BMI, waist-to-hip ratio, HOMA-IR, fat and muscle percentage) was associated with decreased brain glucose metabolism in thalamus, pallidum, hippocampus, putamen and parietal lobe while this cluster was associated with increased brain glucose metabolism in the frontal lobe. Although underlying causal mechanisms are unknown, the results may link with Alzheimer’s dementia. High BMI (47), waist circumference (47), and decreased muscle area (19) are associated with accelerated cognitive decline and a higher probability of Alzheimer’s dementia. Additionally, patient with Alzheimer’s dementia have lower brain glucose metabolism in parietal, temporal lobes (48). A possible explanation is the effect of obesity-driven low-grade systemic inflammation which influences the brain through cytokines (49).

There was a regionally specific effect of glucose levels on brain glucose metabolism. The cluster reflecting both the short-(fasting plasma glucose) and long-term (HbA1c) glycemic levels was negatively associated with brain glucose metabolism in parietal lobe, cingulate, occipital lobe and frontal lobe, whereas positive effects were observed with that in hippocampus, caudate, cerebellum and temporal lobe. Previously, the association of glucose with decreased brain glucose metabolism in parietal lobe, posterior cingulate, prefrontal region and occipital lobe were reported, reflecting a characteristic pattern of Alzheimer’s dementia (23; 50; 51) and possibly through neuronal injury even before the onset of cognitive impairment (52). In addition, FDG uptake competes with endogenous blood glucose and there is region-specific expression of glucose transporters and hexokinase which predominantly affect FDG uptake (53). Also, long-term glucose levels may have a distinct effect on brain glucose metabolism, as short-term glucose level consistently showed the effect on decrease of brain glucose metabolism in regions related with Alzheimer’s dementia. However, the underlying mechanism of this region-based difference remains unclear.

There were several reports regarding stress with brain glucose metabolism of amygdala, insula, striatum (54; 55) and depression with that of hippocampus, frontal lobe (56). The association of psychological cluster with brain glucose metabolism of thalamus and hippocampus became prominent in 5-year follow-up, not in baseline study. Thus, depression and stress may begin to affect brain glucose metabolism after mid 40s when the allostatic load from these psychological cluster has accumulated for long enough.

There was no effect of blood pressure on brain glucose metabolism. However, heart rate had an association with brain glucose metabolism of thalamus, hippocampus, cingulate, cerebellum and occipital lobe. Previously, heart rate was positively associated with brain glucose metabolism of frontal lobe and cingulate (57) and with cerebral blood flow of thalamus, putamen, cerebellum and insula (58). Cardiovascular reactivity to stress might be associated with brain glucose metabolism of cingulate, consistent with previous study with fMRI (59). Also, thalamus and cerebellum influence cardiovascular response by stimulating and inhibiting sympathetic and parasympathetic tone (60; 61).

### Limitations

Only males were included in this study, thus these results may not directly generalize to females. This retrospective study was based on health check-up program. As a brain magnetic resonance imaging (MRI) was not included in the program, MRI-based coregistration and partial volume correction of PET scans could not be done and the results could not be compared with MR-based indices of atrophy. Finally, we did not perform an extensive neuropsychological test on participants so we could not link the changes in regional glucose uptake with cognitive functioning.

## CONCLUSION

Regionally selective decline in brain glucose utilization begins already in the middle age, while individual differences in brain glucose metabolism remain stable. In addition to ageing, brain glucose utilization is also associated with metabolic cluster, blood glucose levels and heart rate. These effects are also consistent over the studied period of 5 years in the middle adulthood.

## DATA AVAILABILITY

The datasets generated during and/or analysed during the current study are available from the corresponding author on reasonable request.

